# Positional Context of Myonuclear Transcription during Injury-induced Muscle Regeneration

**DOI:** 10.1101/2022.02.11.480101

**Authors:** Kole H. Buckley, Andrea L. Nestor-Kalinoski, Francis X. Pizza

**Affiliations:** School of Exercise and Rehabilitation Sciences, University of Toledo, Toledo, Ohio; Advanced Microscopy & Imaging Center, University of Toledo, Toledo, Ohio

## Abstract

Fundamental aspects underlying skeletal muscle regeneration after injury are poorly understood. This investigation begins to address deficiencies in knowledge by examining the kinetics of myonuclear accretion, positioning, and global transcription during injury-induced muscle regeneration in mice. We demonstrate that myonuclear accretion plateaus within 7 d of an injury and that the majority (~70%) of myonuclei are centrally aligned in linear arrays (nuclear chains) throughout the course of regeneration. Relatively few myonuclei were found in a peripheral position (~20%) or clustered (~10%) together during regeneration. Importantly, transcriptional activity of individual myonuclei in nuclear chains was high, and greater than that of peripheral or clustered myonuclei. Transcription occurring primarily in nuclear chains elevated the collective transcriptional activity of regenerating myofibers during the later stage of regeneration. Importantly, the number of myonuclei in chains and their transcriptional activity were statistically correlated with an increase in myofiber size during regeneration. Our findings demonstrate the positional context of transcription during regeneration and highlight the importance of centralized nuclear chains in facilitating hypertrophy of regenerating myofibers after injury.

## Introduction

Skeletal muscle injury resulting from physical activity, trauma, or disease initiates processes that restore structure and function to the injured muscle. These processes are referred to as muscle regeneration and commence with the proliferation of muscle stem cells called satellite cells (Relaix and Zammit, 2012; Dumont et al., 2015). Sustained fusion of progenitor cells derived from satellite cells results in the accumulation of hundreds of nuclei (myonuclei) within newly formed (regenerating) myofibers (Krauss, 2010; Pavlath, 2011; Millay et al., 2014; Bi et al., 2018; Martin et al., 2020). Myonuclear accretion increases the capacity of regenerating myofibers to transcribe genes; which in theory optimizes their maturation through mechanisms that govern transcription and translation. Little however is known about the transcriptional activity of individual myonuclei (Newlands et al., 1998; Kim et al., 2020) and the collective transcriptional activity of myofibers during regeneration. These deficiencies in knowledge limit understanding of how the structure and function of skeletal muscle is restored after injury.

Normally, almost all of the myonuclei within a myofiber are non-randomly dispersed and positioned in a peripheral location near the sarcolemma (Bruusgaard et al., 2003; Bruusgaard et al., 2006). These myonuclei are thought to be positioned in a manner that spatially optimizes molecular and cellular processes of myofiber homeostasis. During regeneration, myonuclei normally align in long centralized linear arrays (nuclear chains) (Clark, 1946; Wada et al., 2008; Martin et al., 2020). Nuclear chains are evident in transverse sections of skeletal muscle by the central position of myonuclei within myofibers, which is a hallmark sign of regenerating myofibers. Regenerating myofibers also have myonuclei situated in non-linear groupings (nuclear clusters) throughout their cytoplasm, as well as in a peripheral position (Clark, 1946; Wada et al., 2008; Liu et al., 2020; Martin et al., 2020). The relative distribution of myonuclei in chains, clusters, and a peripheral position during regeneration has yet to be established. How and why myonuclei are situated in distinctly different positions in regenerating myofibers is unknown.

It is conceivable that the normal positioning of myonuclei in regenerating myofibers facilitates their maturation by spatially optimizing molecular and cellular processes of protein synthesis. This premise is based on several observations. One, prior studies have reported that gene transcripts in developing myotubes/myofibers are translated near myonuclei responsible for their synthesis (Pavlath et al., 1989; Ralston and Hall, 1992) and that proteins within myofibers have limited diffusion (Papadopoulos et al., 2000). Two, a positional context of transcription and translation has been established for myonuclei that normally cluster near the myotendinous and neuromuscular junctions (Hall and Sanes, 1993; Kim et al., 2020; Petrany et al., 2020). These myonuclei have a molecular signature that reflects the unique function of the junction. Lastly, myonuclei in regenerating myofibers do not appear to be transcriptionally equivalent (Newlands et al., 1998; Kim et al., 2020; McKellar et al., 2021). Specifically, prior studies have reported heterogeneity in gene transcripts amongst myonuclei isolated from regenerating muscle (Kim et al., 2020) and qualitative observations of transcriptional diversity amongst myonuclei positioned in nuclear chains during regeneration (Newlands et al., 1998). The extent to which transcriptional activity of myonuclei varies in a manner that reflects their position in regenerating myofibers remains to be determined. We hypothesize that heterogeneity in global transcription exists between myonuclear positions, and that such diversity influences the expansion of myofiber volume (myofiber hypertrophy) that normally occurs during regeneration.

Our primary objective was to examine transcriptional activity of myonuclei and its positional context within regenerating myofibers. Secondarily, we sought to gain insight into the contribution of myonuclear accretion and transcriptional activity to myofiber hypertrophy during regeneration. These objectives were achieved by quantifying myonuclear number, positioning, and global transcription in individual myofibers during injury-induced muscle regeneration in mice.

## Material and Methods

### Mice

Adult (12-16 weeks of age) male and female wild type (C57BL/6) mice were used in this study. Mice were housed and bred in an animal facility at the University of Toledo, which is accredited by the Association for the Assessment and Accreditation of Laboratory Animal Care. Mice were housed with a 12-hour light-dark cycle and fed standard laboratory chow and water ad libitum. Mice were anesthetized using 2.5% isoflurane for surgical procedures and were sacrificed via cervical dislocation prior to muscle collections. All procedures were approved by the institutional animal care and use committee.

### Muscle Injury

Injury to gastrocnemius muscles was achieved via bilateral intramuscular injection of 1.2% barium chloride (BaCl_2_; Sigma-Aldrich) (Hardy et al., 2016; Morton et al., 2019). The gastrocnemius muscle was chosen for this study due to the ability to isolate live myofibers that can be processed for a variety of downstream analyses (e.g., transcriptional activity). The mouse gastrocnemius is composed of primarily (94%) Type II myofibers, which is similar to the commonly used tibialis anterior (Augusto et al., 2017). Muscles were exposed through a small skin incision and a total of 50 μl of BaCl_2_ was injected using a 25-gauge needle and a Hamilton syringe, with each head of the gastrocnemius receiving 25 μl. Skin incisions were sutured closed. Muscles were collected at 7, 14, and 28 days post-injury, as well as from control mice that did not undergo surgery (0 days post-injury).

### Myofiber Isolation and Fixation

Single myofibers were isolated according to published procedures (Au - Pasut et al., 2013; Keire et al., 2013). Briefly, gastrocnemius muscles were enzymatically digested in a well containing 0.18% collagenase type I in Dulbecco modified eagles medium (DMEM) (Sigma-Aldrich) for ~2.5 hours. Isolated myofibers were transferred to wells containing DMEM until a desired number of myofibers were obtained. Myofibers, in a small volume of DMEM, were then transferred to a tube containing 4% paraformaldehyde in phosphate buffered saline (PBS) and incubated for 30 minutes at room temperature. Fixed myofibers were rinsed in PBS and then carefully placed on slides coated with a solution containing 0.04% chromium (III) potassium-sulfate and 0.4% gelatin.

### Detection of Nascent RNA in Myonuclei

Nascent RNA was detected through the use of 5-ethynyluridine (EU) (Invitrogen), a uridine analog that can be incorporated into RNA during transcription (Jao and Salic, 2008). The incorporation of EU into RNA is a global measure of transcription and is not specific to a type of RNA (e.g., mRNA and rRNA). Mice received 2 mg of EU in sterile PBS via i.p. injection 5 hours before muscle collection (Kirby et al., 2016).

Detection of EU within myofibers was achieved using reagents and procedures in the Click-iT^™^ RNA Imaging Kit (Invitrogen). Non-specific/background detection of EU was revealed by omitting the catalyst for the copper mediated reaction from the procedures, as suggested by the manufacturer. Myofibers were treated with DRAQ-5 (1:1000; Invitrogen) to stain myonuclei and then mounted in Fluoromount G (SouthernBiotech).

### Image Acquisition

Myofibers were viewed and imaged using a Leica TCS SP5 multiphoton confocal microscope. Normal (non-regenerating) myofibers from control (0 days post-injury) muscles were imaged. Both regenerating and non-regenerating myofibers were observed in isolates from muscles collected after injury and only regenerating myofibers were imaged. The same settings (e.g., exposure time) were used for imaging of control and regenerating myofibers. The entire length and depth of a myofiber was imaged in 2 μm increments to create a z-stack. A z-projection was produced by merging all images in a single z-stack.

### Quantification of Myonuclei and Transcriptional Activity

Myonuclear number and EU were quantified in z-projections using Image Pro 7 (Media Cybernetics). First, an outline was created of individual nuclei and nuclei were separated from each other via watershed split. The outline was overlaid onto the image of EU. To identify any nuclei in contact with myofibers or outside their boundary, the intensity of EU was temporally increased to exacerbate background fluorescence. Any nuclei in contact with myofibers or outside their boundary were excluded from the analysis.

The total number of myonuclei, as well as the number of myonuclei in chains, clusters, and a peripheral location were counted. For instances in which watershed split was unable to separate individual myonuclei in a close grouping, a built-in algorithm (cluster function) was used to estimate the number of myonuclei in the grouping. A nuclear chain was defined as a series of 5 or more myonuclei that were organized in a linear array near the center of myofibers. A nuclear cluster was defined as a non-linear grouping of 3 or more myonuclei. Lastly, peripheral myonuclei were defined as a myonucleus not localized to a chain or cluster. Myonuclei positioned near the ends of myofibers were not analyzed.

The mean fluorescent intensity (MFI) of EU within outlines of individual myonuclei, as well as the area (μm^2^) of outlines were quantified. This included myonuclei of myofibers that were used to detect background EU. These values were used to calculate the corrected integrated density of EU for each myonucleus using the following equation: (*Myonuclear area* × *MFI*) — (*Myonuclear area* × *MFI* — *background*). The corrected integrated density of EU within individual myonuclei (integrated density/myonucleus) was used to represent their transcriptional activity.

The length and average width of the segment of a myofiber that was used in the quantification of myonuclear number and EU were measured using cellSens software (Olympus Life Sciences). The average width served as a measure of myofiber size and was used in the calculation of myofiber volume (*Myofiber volume* = *π* × *average radius*^2^ × *length of myofiber segment*).

At least 4 myofibers from 3 different muscles per time point were analyzed (n = 14-17 myofibers/time point). On average, 3484 μm of myofiber length (SD = ± 861) and 573 myonuclei per myofiber (SD = ± 253) were analyzed (n = 61 myofibers). The total number of myonuclei analyzed was 34,954.

### Statistics

Data sets were analyzed using one-way or two-way analysis of variance (ANOVA) using Sigma Plot software (Systat). The number of days after injury and myonuclear position were used as grouping factors. The Newman-Keuls *post-hoc* test was used to locate differences when the observed F ratio was statistically significant (p<0.05). To determine relationships between dependent measures, bivariate linear and forward stepwise multiple regression analyses were performed using Sigma Plot. Data is reported as Mean ± SEM.

## Results

### Myonuclear Number and Density

To substantiate and extend our prior work (Martin et al., 2020), we quantified myonuclear number before and during regeneration (Figure 1A). The number of myonuclei in regenerating myofibers (myonuclei/100 μm) was 63-73% higher at 7, 14, and 28 days post-injury compared to control (0 days post-injury) levels (Figure 1B). This is consistent with our previous findings that myonuclear number in regenerating myofibers exceeds that of non-regenerating myofibers within control muscle (Martin et al., 2020). Importantly, myonuclear number in regenerating myofibers was similar at 7, 14, and 28 days post-injury. This finding indicates that myonuclear accretion reaches a plateau during an early stage of regeneration.

**Figure 1.**
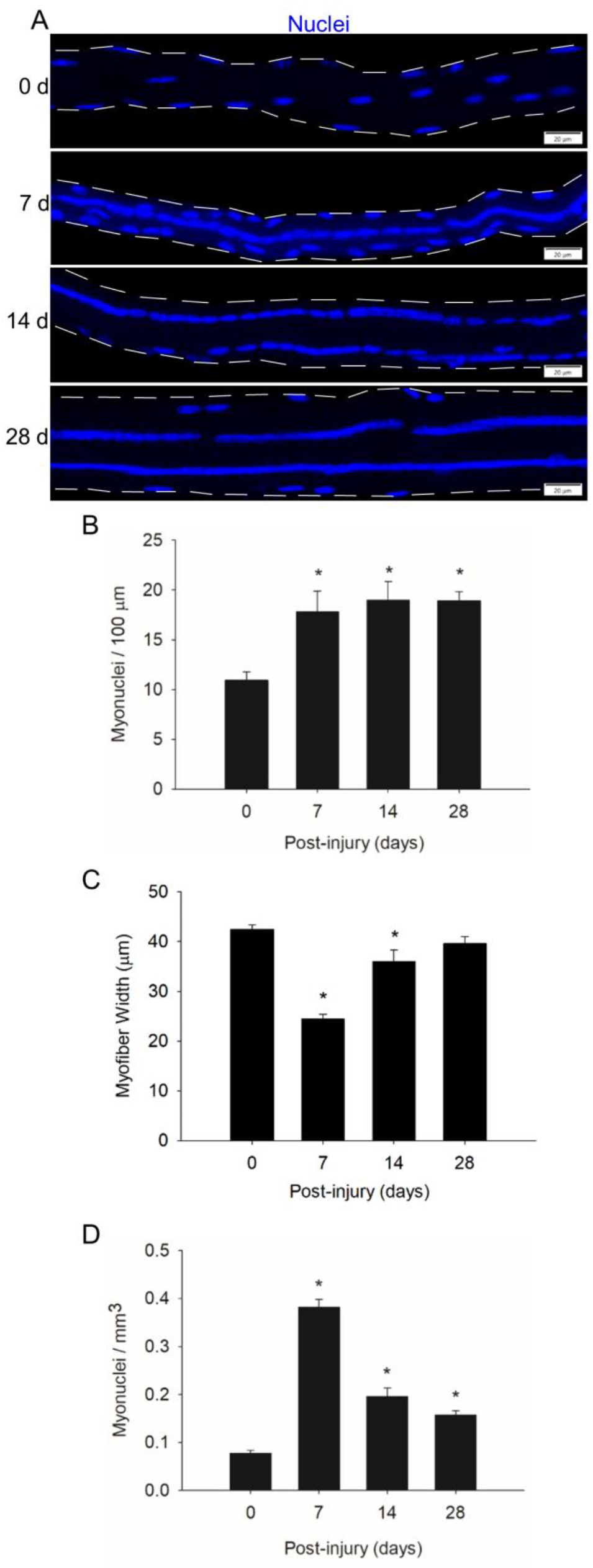
Myonuclear number and density before and during injury-induced muscle regeneration. (**A**) z-projection images of myonuclei (blue) in myofibers isolated at 0, 7, 14, and 28 d post-injury. Scale bar = 20 μm. (**B**) The number of myonuclei expressed relative to myofiber length (100 μm). (**C**) The average width (μm) of myofibers. (**D**) The number of myonuclei expressed relative to myofiber volume (mm^3^). n = 14-17 myofibers per time point. * = Different at indicated time point compared to 0 d post-injury (p < 0.001).

To examine myonuclear density during regeneration, we expressed myonuclear number relative to myofiber volume (myonuclei/mm^3^). Myofiber width, which was used in calculating myofiber volume, was 42% lower at 7 days post-injury compared to control levels and progressively increased during prolonged recovery (Figure 1C). These findings are consistent with the progressive increase in myofiber cross-sectional area, which is indicative of hypertrophy, during the course of regeneration (Martin et al., 2020).

Myonuclear density increased by 4.9-fold at 7 days post-injury and steadily decreased from 7 to 28 days post-injury (Figure 1D). The decrease in myonuclear density during the course of regeneration likely reflects the expansion of myofiber volume, as opposed to the loss of myonuclei. This interpretation is based on the finding that myofibers were undergoing hypertrophy during regeneration (Figure 1C) while myonuclear number (myonuclei/100μm) remains relatively constant (Figure 1B) (Martin et al., 2020). Furthermore, we are not aware of any evidence of myonuclear apoptosis or autophagy during regeneration.

### Myonuclear Positioning

We quantified the number of myonuclei situated in nuclear chains and clusters, as well as in their normal peripheral position (Figure 2A). This was done to better characterize myonuclear positioning during regeneration. Nuclear clusters that normally form near the myotendinous junction (Bruusgaard et al., 2003; Liu et al., 2020) were not included in our analysis as we did not analyze the ends of myofibers. On the other hand, we analyzed the middle region of myofibers, which would include myonuclei that normally cluster in the neuromuscular junction (Bruusgaard et al., 2003; Grady et al., 2005).

**Figure 2.**
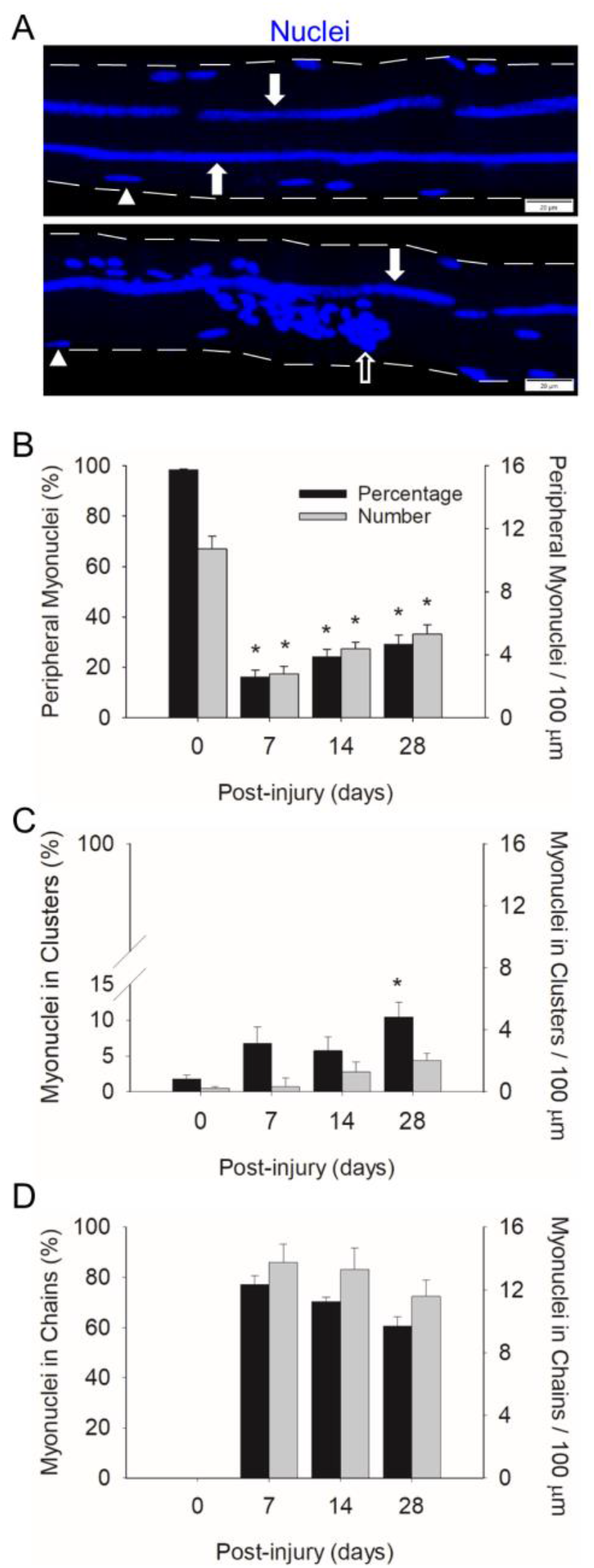
Myonuclear positioning before and during injury-induced muscle regeneration. (**A**) z-projection images of myonuclei (blue) positioned in chains (filled arrow), clusters (open arrow), and a peripheral position (arrowhead) during regeneration. Scale bar = 20 μm. The number of myonuclei in a peripheral position (**B**), clusters (**C**), and chains (**D**) expressed as a percentage of the total number myonuclei within a myofiber (Black bars) and relative to 100 μm of myofiber length (grey bars). n = 14-17 myofibers per time point. * = Different at indicated time point compared to 0 d post-injury (p < 0.001 for peripheral myonuclei and p < 0.05 for clustered myonuclei).

In control myofibers (0 days post-injury), 98% of the myonuclei were in a peripheral position (Figure 2B). The remaining 2% of myonuclei resided in nuclear clusters (Figure 2C). These clusters likely represent myonuclei near the neuromuscular junction. As expected, no nuclear chains were observed in control myofibers (Figure 2D).

Peripheral myonuclei were substantially reduced during regeneration. Specifically, the number of peripheral myonuclei, expressed relative to myofiber length, was 74% lower at 7 compared to 0 days post-injury. Both the number and percentage of peripherally located myonuclei increased gradually from 7 to 28 days post-injury. Importantly, the majority of myonuclei during regeneration resided in centralized nuclear chains. Indeed, 77% of the myonuclei were situated in chains at 7 days post-injury. This number progressively decreased to 60% by 28 days post injury. The number of clustered myonuclei, expressed relative to myofiber length, tended (p = 0.053) to increase during regeneration; whereas the percentage of myonuclei in clusters increased to 10% at 28 d post-injury.

Our findings demonstrate a striking change in the position of myonuclei during regeneration. Myonuclei in regenerating myofibers are primarily situated in centralized nuclear chains during regeneration, which is in mark contrast to the peripheral position of myonuclei in control myofibers.

### Myofiber Size is Responsive to Myonuclear Number and Position

To explore the premise that myonuclear accretion promotes myofiber hypertrophy during regeneration, we performed correlational analyses using myonuclear number and myofiber volume (mm^3^). When all post-injury time points were included in our analysis, a strong positive correlation was observed for regenerating myofibers (r = 0.77; Figure 3A). This finding is consistent with the paradigm that myonuclear accretion facilitates myofiber hypertrophy during regeneration.

**Figure 3.**
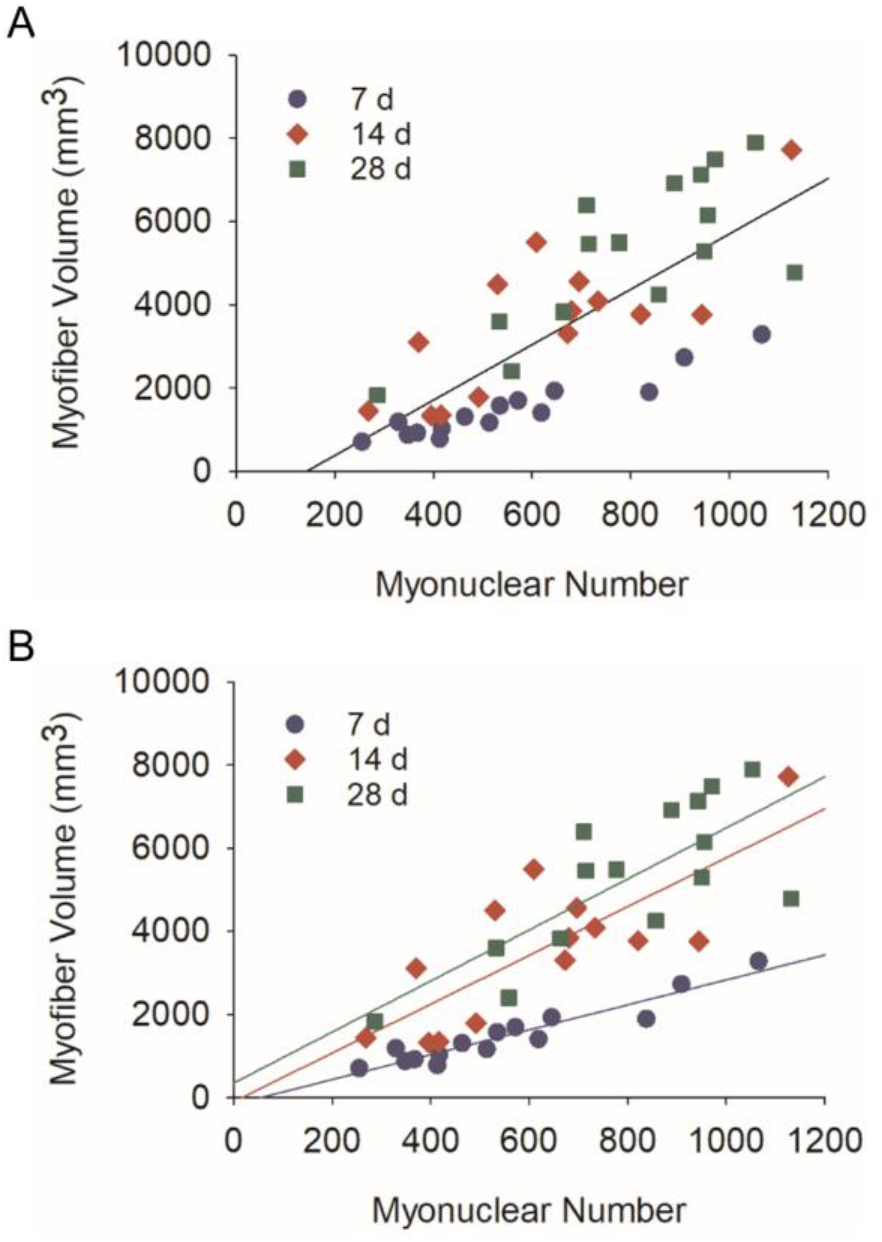
Relationship between myonuclear number and myofiber volume during regeneration. Scatter plots and regression lines are shown for the number of myonuclei within a segment of a regenerating myofiber and its corresponding volume (mm^3^). (**A**) Data and regression line (r = 0.77) for 7, 14, and 28 days post-injury (Myofiber volume (mm^3^) = 6.87×number of myonuclei – 1072.8; n = 44 myofibers). (**B**) Data and regression lines for 7 days (r = 0.95; Myofiber volume (mm^3^) = 2.99×number of myonuclei – 158.9; n = 15 myofibers), 14 days (r = 0.78; Myofiber volume (mm^3^) = 5.86×number of myonuclei – 94.2; n = 14 myofibers), and 28 days (r = 0.82; Myofiber volume (mm^3^) = 6.96×number of myonuclei −244.1; n = 15 myofibers) post-injury. All correlation coefficients were statistically significant (p < 0.01).

Strikingly, a near perfect correlation was observed between myonuclear number and myofiber volume at 7 days post-injury (r = 0.95; Figure 3B). Compared to 7 d post-injury, correlation coefficients were lower at 14 (r = 0.78) and 28 (r = 0.82) days post-injury. In contrast, the slope of the regression line increased after 7 days post-injury. These findings indicate that the size of regenerating myofibers is closely coupled to myonuclear number during the early stage of regeneration. They also indicate that regenerating myofiber size becomes more responsive to myonuclei after myonuclear accretion has plateaued (Figure 1B). Increased transcriptional activity and/or translation could explain the enhanced responsiveness of regenerating myofibers to myonuclei during the later stage of regeneration.

We also examined the relationship between myonuclear positioning and myofiber size during regeneration using stepwise multiple regression analysis. This analysis was performed on data sets that included all post-injury time points, as well as from each post-injury time point (Table 1). Overall, these analyzes revealed that the number of myonuclei in nuclear chains is a strong predictor of myofiber volume. The inclusion of myonuclei in a peripheral location strengthened the prediction of myofiber volume at 28 d post-injury. In contrast, the inclusion of myonuclei in clusters did not enhance the prediction of myofiber volume at any time point. These findings indicate that myonuclear positioning influences myofiber hypertrophy after injury. Specifically, myonuclei situated in nuclear chains appear to be important in facilitating hypertrophy of regenerating myofibers.

**Table 1.** Relationship Between Myonuclear Position and Myofiber Volume

| 7-28 d Multiple R = 0.83 |  |  |  |
| --- | --- | --- | --- |
| Myonuclear Position | Slope | Std. Error | P-value |
| Chains | 6.354 | 0.969 | <0.001 |
| Peripheral | 14.395 | 2.111 | <0.001 |
| Clusters | - | - | N.S. |
| Intercept | -1559.436 | 556.026 | - |
| 7 d Multiple R = 0.95 |  |  |  |
| Chains | 3.326 | 0.257 | <0.001 |
| Peripheral | - | - | N.S. |
| Clusters | - | - | N.S. |
| Intercept | -79.108 | 141.003 | - |
| 14 d Multiple R = 0.77 |  |  |  |
| Chains | 7.230 | 1.725 | 0.001 |
| Peripheral | - | - | N.S. |
| Clusters | - | - | N.S. |
| Intercept | 358.713 | 824.064 | - |
| 28 d Multiple R = 0.92 |  |  |  |
| Chains | 8.994 | 1.186 | <0.001 |
| Peripheral | 11.997 | 2.396 | <0.001 |
| Clusters | - | - | N.S. |
| Intercept | -1692.868 | 934.934 | - |

### Transcriptional Activity of Myonuclei

In theory, elevated transcription is needed to drive increases in myofiber protein synthesis and size during regeneration. Thus, we investigated the transcriptional activity of individual myonuclei during regeneration by quantifying EU in individual myonuclei (Figure 4). The integrated density of EU within individual myonuclei was used to represent their transcriptional activity.

**Figure 4.**
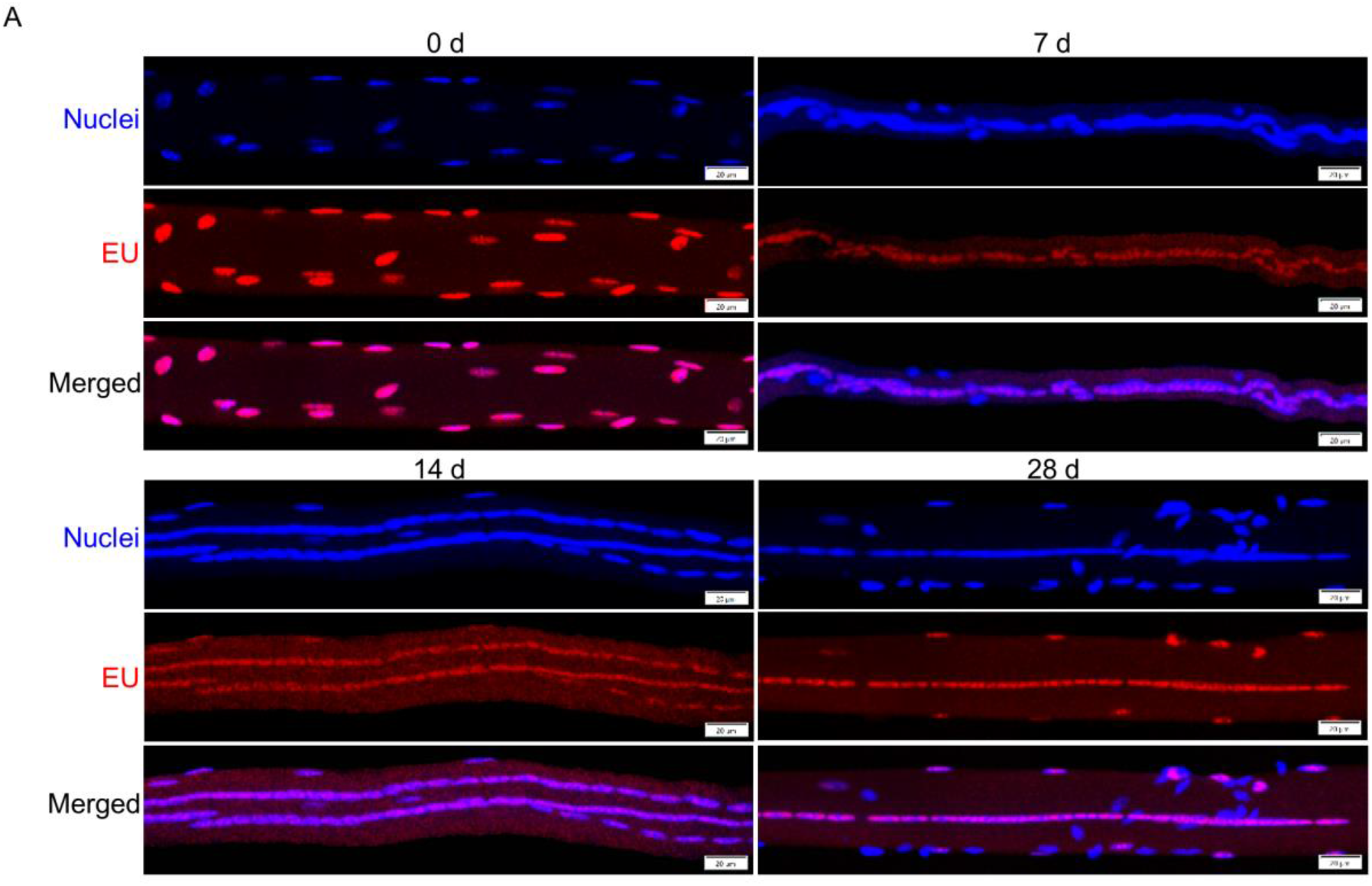
Detection of nascent RNA in myofibers before and during injury-induced muscle regeneration. Mice were administered 5-ethynylurdine (EU; red) and its presence within myonuclei (blue) was detected as described in Methods. Images are representative of responses observed at 0, 7, 14, and 28 d post-injury. Scale bar = 20 μm.

Unexpectedly, the mean integrated density for individual myonuclei in regenerating myofibers was 16-30% lower than that observed in control myofibers (Figure 5A). These findings demonstrate an overall suppression in the transcriptional activity of individual myonuclei during regeneration.

**Figure 5.**
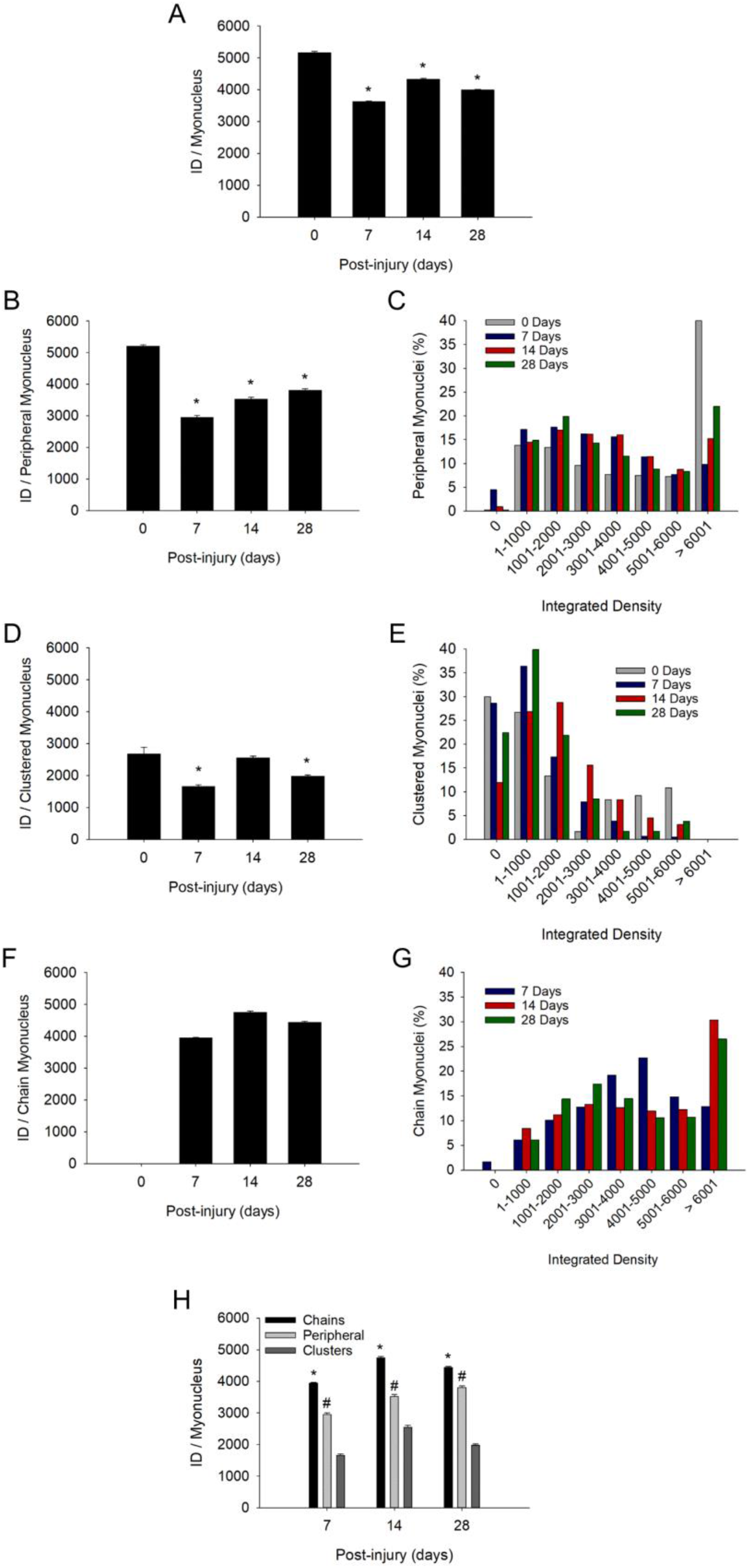
Transcriptional activity of individual myonuclei before and during injury-induced muscle regeneration. Transcriptional activity of individual myonuclei was determined by calculating the integrated density (ID) of EU for each myonucleus as described in Methods. (**A**) ID for all myonuclei, independent of their position within a myofiber (n = 6,049 - 11,857 myonuclei/time point). (**B**) ID for myonuclei in a peripheral position (n = 1,275 – 3,230) myonuclei/time point). (**C**) Frequency distribution of integrated densities for peripheral myonuclei. Data is expressed as a percentage of the total number of peripheral myonuclei at each time point. (**D**) ID for myonuclei in nuclear clusters (n = 120 – 1,337 myonuclei/time point). (**E**) Frequency distribution of integrated densities for myonuclei in nuclear clusters expressed as a percentage of the total number of clustered myonuclei at each time point. (**F**) ID for myonuclei in nuclear chains (n = 6,424 – 7,290 myonuclei/time point). (**G**) Frequency distribution of integrated densities for myonuclei in nuclear chains. Data is expressed as a percentage of the total number of myonuclei situated in nuclear chains at each time point. * in (**A**), (**B**), (**D**), and (**F**) = lower at indicated time point compared to 0 days post-injury (p < 0.001). (**H**) ID for myonuclei in each myonuclear position at 7, 14, and 28 days post-injury (n = 120 - 11,857 myonuclei/myonuclear position and time point). * = higher for myonuclei in nuclear chains compared to the other positions (p < 0.001). # = higher for myonuclei in a peripheral position compared to those in nuclear clusters (p < 0.001).

### Positional Context of Transcriptional Activity of Myonuclei

A major goal was to determine the extent to which transcriptional activity of myonuclei varies in a manner that reflects their position in regenerating myofibers. Thus, we determined if the reduced transcriptional activity of myonuclei (Figure 5A) was specific to a myonuclear position. This was done by segregating the integrated density of individual myonuclei into three myonuclear positions.

During regeneration, the mean integrated density for individual myonuclei in a peripheral location was reduced by 43, 32, and 27% at 7, 14, and 28 days post-injury, respectively (Figure 5B). Many peripheral myonuclei at 0 d post-injury had very high transcriptional activity (integrated density > 6000) (Figure 5C). This population of peripheral myonuclei was substantially reduced at 7 days post-injury and then gradually increased during the remaining days of recovery. The mean integrated density for clustered myonuclei was reduced by 5-38% during regeneration (Figure 5D). A large percentage of the myonuclei in nuclear clusters had very low transcriptional activity (integrated density = 0-1000) before and during regeneration (Figure 5E).

Importantly, reductions in the transcriptional activity of peripheral and clustered myonuclei during regeneration were accompanied by the emergence of myonuclei in nuclear chains (Figure 2D) and their high transcriptional activity (Figure 5F). The mean integrated density for individual myonuclei in nuclear chains remained high during the course of regeneration and reached levels that were 76-91% of the mean for peripheral myonuclei at 0 days post-injury. The percentage of myonuclei in nuclear chains that had very high transcriptional activity (integrated density > 6000) increased substantially at 14 days post-injury and remained high at 28 days post-injury (Figure 5G).

Population specific differences in transcriptional activity were also apparent when comparing transcriptional activity between myonuclear positions during regeneration (Figure 5H). The mean integrated density for individual myonuclei in nuclear chains was 22% and 53% higher than that observed for peripheral and clustered myonuclei, respectively. Furthermore, peripheral myonuclei had a 40% greater mean integrated density compared to myonuclei in nuclear clusters.

Our findings demonstrate for the first time the positional context of transcription in regenerating myofibers. Myonuclei in nuclear chains were transcriptionally very active throughout the course of regeneration. In contrast, the transcriptional activity of peripheral and clustered myonuclei were suppressed during regeneration. Heterogeneity in transcriptional activity of individual myonuclei was also apparent within each myonuclear position, as indicated in frequency distribution plots.

### Transcriptional Activity of Myofibers

Conceptually, the collective transcriptional activity of a myofiber reflects both the number of myonuclei within them, and the transcriptional activity of each myonucleus. Because regenerating myofibers had more myonuclei (Figure 1B) with reduced activity (Figure 5A) compared to control myofibers, we examined the collective transcriptional activity within myofibers. This was done by summing the integrated density for all myonuclei within a myofiber and then expressing the total integrated density relative to myofiber length (integrated density/ μm) or volume (integrated density/mm^3^).

Total integrated density per myofiber length (integrated density/μm) was 49 and 39% higher than control levels at 14 and 28 days post-injury, respectively (Figure 6A). This indicates that the reduced transcriptional activity of individual myonuclei during regeneration (Figure 5A) was offset by an elevated number of myonuclei within regenerating myofibers (Figure 1B). Thus, the collective transcriptional activity of a regenerating myofiber exceeded that of a control myofiber.

**Figure 6.**
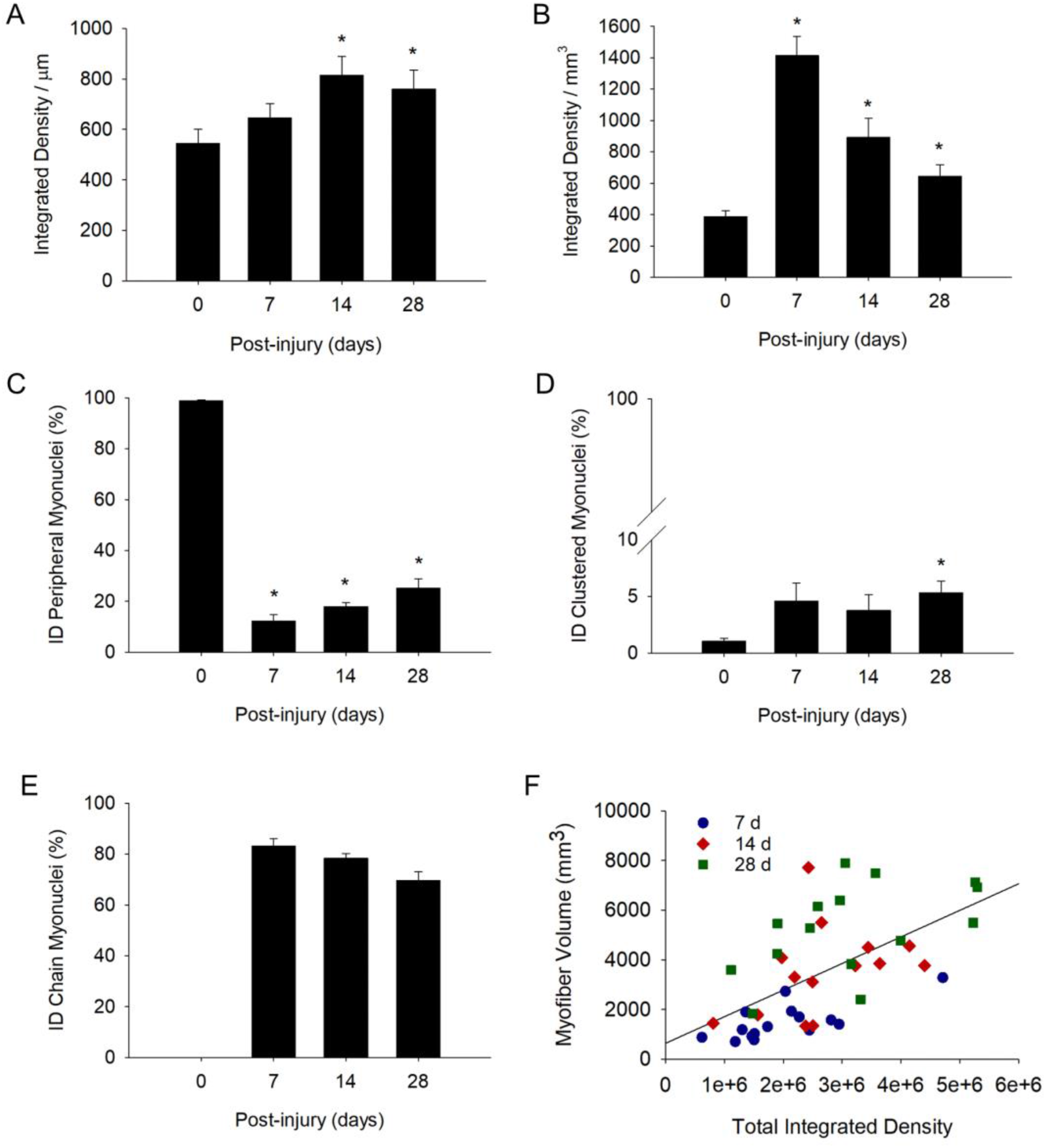
Transcriptional activity of myofibers before and during injury-induced muscle regeneration. Transcriptional activity of myofibers was determined by expressing the total integrated density within a myofiber relative to myofiber length (integrated density/μm; (**A**) or volume (integrated density/mm^3^; (**B**). * = Different at indicated time point compared to 0 d post-injury (p < 0.05 in (**A**) and p < 0.001 in (**B**)). We also expressed total integrated density (ID) for peripheral (**C**), clustered (**D**), and chain (**E**) myonuclei as a percentage of the total integrated density within the myofiber (n = 14-17 myofibers per time point). * = Different at indicated time point compared to 0 d post-injury (p < 0.001 for peripheral myonuclei and p < 0.05 for clustered myonuclei). (**F**) Scatter plot and regression line (r = 0.59) for the total integrated density (independent of myonuclear position) within a segment of a regenerating myofiber and its corresponding volume (mm^3^). Plot includes data at 7, 14, and 28 days post-injury (Myofiber volume (mm^3^) = 634.8×total integrated density + 0.001; n = 44 myofibers).

Total integrated density expressed relative to myofiber volume (integrated density/mm^3^) was also higher after injury (Figure 6B). Specifically, integrated density/mm^3^ was 3.7, 2.3, and 1.7 fold higher than control levels at 7, 14, and 28 days post-injury, respectively. The progressive decline in integrated density/mm^3^ most likely reflects the expansion of myofiber volume. This is because both the number of myonuclei (myonuclei/100um; Figure 1B) and transcriptional activity of individual myonuclei (integrated density/myonuclei; Figure 5A) were relatively constant during regeneration.

### Positional Context of Transcription in Myofibers

We next examined the relative contribution of each myonuclear position to the collective transcriptional activity of myofibers. This was done by summing integrated densities for each myonuclear position within a myofiber and then expressing the total integrated density for each position as a percentage of the total integrated density within the myofiber.

In control myofibers, peripherally positioned myonuclei accounted for 99% of the total integrated density (Figure 6C), while clusters made up the other 1% (Figure 6D). During regeneration, 83% of the total integrated density at 7 days post-injury was attributable to myonuclei situated in nuclear chains (Figure 6E). The contribution of nuclear chains to the total integrated density within a myofiber declined to 69% at 28 days post-injury. Meanwhile, the contribution of peripheral myonuclei to the total integrated density was low initially (12% at 7 days post-injury), and progressively increased to 25% at 28 days post-injury. Myonuclei in nuclear clusters contributed very little (3-5%) to the total integrated density within a myofiber during regeneration.

These findings demonstrate that most of the transcription in a regenerating myofiber is occurring within nuclear chains. Their contribution diminishes, while the contribution of peripheral myonuclei to transcription increases over the course of regeneration.

### Myofiber Size is Responsive to the Positional Context of Transcription

As elevations in transcription is predicted to promote myofiber hypertrophy during regeneration, we examined the relationship between total integrated density and myofiber volume. A moderate correlation (r = 0.59; Figure 7A) was observed between total integrated density and myofiber volume when all post-injury time points were analyzed. The correlation coefficient was higher at 7 days post-injury (r =0.72) and lower at both 14 (r = 0.38) and 28 (r = 0.50) days post-injury. Overall, our findings are consistent with the paradigm that myonuclear transcription facilitates myofiber hypertrophy during regeneration.

We also examined the relationship between the positional context of transcription and myofiber volume using stepwise multiple regression analysis (Table 2). These analyzes revealed that the collective transcriptional activity of myonuclei in nuclear chains is a predictor of myofiber volume during regeneration. The inclusion of the collective transcriptional activity of peripheral and clustered myonuclei did not enhance the prediction of myofiber volume. These findings are consistent with the premise that transcription occurring in nuclear chains facilitates hypertrophy of regenerating myofibers.

**Table 2.** Integrated Density of Myonuclear Positions vs. Myofiber Volume

| **Myonuclear Position** | **Slope** | **Std. Error** | **P-value** |
| --- | --- | --- | --- |
| Chains | 0.00125 | 0.000318 | <0.001 |
| Peripheral | - | - | N.S. |
| Clusters | - | - | N.S. |
| Intercept | 970.977 | 690.304 | - |

## Discussion

Prior work has established that regenerating myofiber formation after injury is dependent on satellite cells (Relaix and Zammit, 2012) and the fusion of progenitor cells derived from them (Millay et al., 2014; Bi et al., 2018). What remains to be established are the processes that govern the maturation of regenerating myofibers into a normal sized myofiber with peripheral myonuclei. Specifically, little is known about the kinetics of myonuclear accretion and positioning, and their impact on transcription and myofiber size during regeneration. The present study addresses these deficiencies in knowledge and reveals novel aspects of muscle regeneration. Namely, we demonstrate heterogeneity in transcriptional activity between myonuclear positions during regeneration. Myonuclei situated in nuclear chains were transcriptionally very active; whereas the transcriptional activity of peripheral and clustered myonuclei was suppressed during regeneration. The large number of myonuclei in nuclear chains and the transcription within them were statistically associated with an increase in myofiber size during regeneration. Taken together, our findings indicate that the positioning and transcriptional activity of myonuclei in nuclear chains facilitates myofiber hypertrophy during regeneration.

We were the first to quantify myonuclear number during injury-induced muscle regeneration (Martin et al., 2020). The present study extends our initial findings by demonstrating that regenerating myofibers obtain the majority of their myonuclei within 7 days of an injury. This interpretation is based on the finding that myonuclear number (myonuclei/100μm) was elevated above control levels at 7 days post-injury and remained relatively constant during prolonged recovery. The observed kinetics in myonuclear accretion is consistent with the robust myogenic cell proliferation and fusion that normally occurs within 7 days of an injury (Millay et al., 2014; Dumont et al., 2015; Hardy et al., 2016; Webster et al., 2016; Bi et al., 2018). Why regenerating myofibers contain a greater number of myonuclei (myonuclei/100μm) than control myofibers is unknown (Martin et al., 2020) (present study). We suspect that the elevated number of myonuclei in regenerating myofibers reflects the pro-fusion environment of injured muscle, as opposed to the involvement of an unknown mechanism that dictates myonuclear number during an early stage of regeneration.

A novel finding of the present study is the dramatic change in the position of myonuclei during regeneration. During regeneration, only ~20% of the myonuclei in regenerating myofibers were found in a peripheral position. This is in mark contrast to control myofibers, which had 98% of their myonuclei in a peripheral location. Most of the myonuclei (~70%) in regenerating myofibers were found in nuclear chains, which are not present in control myofibers. Few (<10%) myonuclei in regenerating myofibers were found in nuclear clusters. Interestingly, myonuclei situated in nuclear chains diminished, while the number of peripheral myonuclei increased during the course of regeneration. This finding may reflect the movement of myonuclei from centralized chains to a peripheral position as the regenerating myofiber matures. This interpretation is consistent with the movement of myonuclei from a central to a peripheral position during muscle development (Harris et al., 1989; White et al., 2010) and in vitro myogenesis (Wilson and Holzbaur, 2015; Espigat-Georger et al., 2016; Roman et al., 2017).

Researchers have begun to identify mechanisms governing myonuclear positioning. Using mammalian, invertebrate (e.g., drosophila melanogaster) and in vitro models, investigators have demonstrated the importance of the LINC complex (Linker of Nucleoskeleton and Cytoskeleton) and motor proteins in the normal peripheral positioning of myonuclei (Folker and Baylies, 2013; Volk, 2013; Roman and Gomes, 2018; Azevedo and Baylies, 2020). Specifically, impairments in the expression of LINC complex or motor proteins during muscle development have been reported to cause myonuclei to cluster or align near the sarcolemma (Zhang et al., 2007; Lei et al., 2009; Zhang et al., 2010; Metzger et al., 2012; Liu et al., 2020). This aberrant myonuclear positioning is distinctly different from the normal positioning of myonuclei within centralized nuclear chains during regeneration. Less is known about the specific contribution of the cytoskeleton, consisting of microtubules, intermediate filaments, and actin filaments, to myonuclear positioning (Folker and Baylies, 2013; Roman and Gomes, 2018; Becker et al., 2020). Interestingly, the directionality of microtubules changes to a more longitudinal orientation during regeneration (Randazzo et al., 2019), which could contribute to the alignment of myonuclei in nuclear chains. Nevertheless, the extent to which the LINC complex, motor proteins, and/or the cytoskeleton mediate myonuclear positioning in regenerating myofibers remains to be determined.

Myonuclear accretion occurs during postnatal development (Moss and Leblond, 1971; White et al., 2010; Bachman et al., 2018; Cramer et al., 2020) and after resistance exercise training (Egner et al., 2016; Goh and Millay, 2017; Murach et al., 2021). In both cases, myonuclei appear to be added to existing myofibers and positioned in a peripheral location. During postnatal development in mice, myonuclear number and myofiber size increase in parallel prior to puberty (White et al., 2010; Bachman et al., 2018). The myonuclear accretion during this period has been statistically and mechanistically linked to myofiber hypertrophy (White et al., 2010; Bachman et al., 2018; Cramer et al., 2020). In the present study, myonuclear number was statistically associated with an increase in myofiber size, suggesting that myonuclear accretion also facilitates hypertrophy of regenerating myofibers. Importantly, the statistical association between myonuclei number and myofiber size was almost exclusively the result of myonuclei situated in nuclear chains. Taken together, our findings indicate that hypertrophy of regenerating myofibers primarily reflects the function of myonuclei situated in nuclear chains.

A major finding of the present study was the heterogeneity in transcriptional activity between and within myonuclear positions during regeneration. Strikingly, the transcriptional activity of peripheral myonuclei in regenerating myofibers was suppressed. This suppression was specific to regenerating myofibers as the transcriptional activity of peripheral myonuclei in non-regenerating myofibers within regenerating muscle was very high (unpublished observations). In contrast, myonuclei situated in nuclear chains were transcriptionally very active during regeneration. The heterogeneity observed in transcriptional activity amongst myonuclei in nuclear chains is consistent with qualitative observations of Newlands et al. (Newlands et al., 1998). Our novel findings indicate that transcription in regenerating myofibers is regulated in a manner that reflects myonuclear positioning.

In contrast to our detailed understanding of transcription in proliferating and differentiating myogenic cells (Parker et al., 2003; Braun and Gautel, 2011), the identity and spatial distribution of regulators of transcription (e.g., transcription factors) in regenerating myofibers remains to be determined. Myonuclei in nuclear chains are very closely aligned in a central location between myofibrils within regenerating myofibers (Newlands et al., 1998; Wada et al., 2008; Martin et al., 2020); whereas those in a peripheral position are separated from each other and near the sarcolemma (Bruusgaard et al., 2003; Bruusgaard et al., 2006). These anatomical differences could explain the positional context of transcription if regulators of transcription tend to localize near nuclear chains and/or if there is an exchange of nuclear content between neighboring myonuclei within nuclear chains. It is also conceivable that the positioning of nuclear chains between myofibrils results in greater mechanotransduction of myonuclei within nuclear chains compared to those in a peripheral position. Further investigation is needed to test the validity of such scenarios.

Myonuclear accretion was accompanied by an elevation in the collective transcriptional activity of regenerating myofibers. This was achieved despite reductions in the number and transcription activity of peripheral myonuclei. Stated differently, the large number of myonuclei in nuclear chains, in conjunction with their high transcriptional activity, elevated transcription in regenerating myofibers to levels that exceeded those found in control myofibers. These findings indicate that the increased demand for transcription during regeneration is met primarily by myonuclei situated in nuclear chains. Transcription occurring in nuclear chains appears to be physiologically relevant as the collective transcriptional activity in nuclear chains was statistically associated with an increase in myofiber size during regeneration. Our findings also indicate that there is a degree of transcriptional coordination amongst myonuclei in nuclear chains and a peripheral location to meet the overall demand for transcription. In essence, there appears to a division of transcriptional labor between myonuclear positions during regeneration. The molecular underpinnings for the division of labor amongst myonuclei in regenerating myofibers remain to be determined.

The technique used in the present study to quantify transcriptional activity is not specific to a type of RNA (Jao and Salic, 2008). Given that most of the RNA in cells is ribosomal RNA (Figueiredo and McCarthy, 2019), our findings likely reflect transcription for components of ribosomes during regeneration. It is also likely that some of the transcriptional activity observed in regenerating myofibers altered messenger RNA levels, and possibly the levels of other types of RNA. Our speculation is supported by recent studies that have demonstrated that the transcriptome in regenerating muscle includes transcripts that promote ribosomal biogenesis and that encode a large number and range of proteins (Dell’Orso et al., 2019; De Micheli et al., 2020; Kim et al., 2020; Oprescu et al., 2020; McKellar et al., 2021). Another limitation of our analysis is that some of the transcription observed in peripheral and clustered myonuclei may reflect transcriptional activity within satellite cells and/or other cells that are very closely associated with the sarcolemma. The observed suppression in global transcription in peripheral and clustered myonuclei during regeneration, however, is in contrast to increased gene expression in satellite cells/myoblasts after injury (Dell’Orso et al., 2019; De Micheli et al., 2020; Oprescu et al., 2020). Nevertheless, our findings warrant investigation into the type of RNA being transcribed during regeneration, as well as the spatial distribution of transcription factors and transcripts in regenerating myofibers. This information will be fundamentally important in understanding how and why myonuclei are situated in distinctly different positions in regenerating myofibers.

The present study provides a deeper understanding of myonuclear number, positioning, and transcriptional activity during injury-induced regeneration. The results presented herein provide a foundation of knowledge for future investigations into the spatial regulation of transcription in regenerating myofibers, as well as downstream mechanisms that facilitate their maturation. Knowledge gained from work in this area will facilitate the development of therapeutic approaches for restoring structure and function to injured or diseased muscle.

## Author Contributions

K.H.B. conceived of the study and designed all experiments, acquired, analyzed, and interpreted all data as well as wrote the manuscript. A.L.N-K. aided in confocal image acquisition. F.X.P. conceived of the study, designed experiments, analyzed and interpreted all data, and wrote the manuscript.

